# Recombinant production and purification of the human protein Tau

**DOI:** 10.1101/545566

**Authors:** Luca Ferrari, Stefan G.D. Rüdiger

## Abstract

Tau protein is a microtubule-stabilizing protein whose aggregation is linked to Alzheimer’s Disease and other forms of dementia. Tau biology is at the heart of cytoskeletal dynamics and neurodegenerative mechanisms, making it a crucial protein to study. Tau purification, however, is challenging as Tau is disordered, which makes it difficult to produce in recombinant system and is degradation-prone. It is thus challenging to obtain pure and stable preparations of Tau. Here, we present a fast and robust protocol to purify Tau recombinantly in Escherichia coli. Our protocol allows purifyig Tau either tag-less free or FLAG-tagged at its N-terminus. By exploiting a cleavable affinity tag and two anion exchange columns, we obtained Tau is of high purity, stable and suitable for in vitro studies, including aggregation experiments that resemble neurodegenerative processes.

## INTRODUCTION

Tau aggregation is the prominent signature of Alzheimer and other diseases termed tauopathies (1–3) It is linked to neuronal death and sufficient to induce neurodegeneration (2, 4). In Alzheimer, aggregation of Tau into fibrils correlates with cognitive impairment in humans (5). Understanding Tau biology is thus of paramount importance to understand neurodegeneration.

Tau protein is predominantly expressed in the central nervous system, with six documented isoforms obtained from alternative splicing and developmentally regulated (4). Tau belongs to the family of microtubule associated proteins, or MAPs, collectively orchestrating microtubule assembly (6), This family of proteins act redundantly, as Tau *null* mice show no phenotype and have increased levels of the paralog MAP1A (7). Studies attempting to propose a role of Tau in axon growth yielded varying results, based on the cell lines investigated (7, 8). Recent data demonstrated how Tau and other MAPs compete to distribute cargos correctly ((9)). Thus, Tau is bound to microtubules in physiological state, and its detachment from them is a necessary step preceding aggregation and pathology.

Tau binding to microtubules is highly dynamic and regulated by phosphorylation, with Tau having more than 40 potential phosphorylation sites (10). Phosphorylation and consequent detachment from microtubule are thought to be the initial steps triggering Tau aggregation in dementia, however recent structural data show that phospho-sites do not have a structural role in the assembly of Tau aggregates (11, 12). Also, hyperphosphorylation is fundamental for hibernation of rodents but does not lead to dementia (6). Finally, some phosphorylation sites may have a protective role against Tau aggregation (13). Thus, Tau hyper-phosphorylation cannot explain alone aggregation and disease progression.

Tau aggregation may be caused by the collapse of the chaperone network instead (14). Tau sequence itself endows the potential for Tau aggregation. Tau is a disordered protein of 441 residues in its longest isoform (4, 15). Tau N-terminus composition enhance its solubility, with few hydrophobic residues (10.6 non hydrophobic residues per hydrophobic one (16)). On the contrary, microtubule binding region has higher hydrophobic content (4.4 non hydrophobic residues per hydrophobic one), and indeed is the region comprising the core of amyloid structures of Alzheimer’s and Pick’s Disease (11, 12). Although Tau is soluble and dynamic, its own sequence has the potential to convert to toxic folds. Surprisingly, it is hard to pinpoint the exact role of aggregates in neurodegeneration (17) Thus, Tau pathology is left with many unresolved issues, making it necessary to develop new approaches to study it. These approaches include generating stable and reliable preparations of Tau. Obtaining pure Tau preparations is however challenging, as Tau disordered nature makes it vulnerable to protease action (15). Tau stability and purity are essential to perform a variety of assays, for instance aggregation experiments recapitulating neurodegeneration. Current available protocols focus on purifying tag-less versions, however this may limit downstream applications relying on Tau recognition by an antibody, such as pull-downs or western blot detection (18, 19).

Here, we present an alternative protocol to obtain recombinant Tau from *Escherichia coli*. Compared to previous protocol, we introduced a cleavable tag His_6_-Smt at the N-terminus to maintain a tag-less version of the protein. We introduced a second anion exchange purification step to remove the tag and to further decrease impurities. Despite this extra purification step, the resulting protocol was still fast, meaning that Tau purification – from cell lysate to pure protein – takes 1 to 2 days, based on the purified variant. We show that the protocol robustly purified also a FLAG-tagged version of Tau at the N-terminus. This variant can be used for instance to perform pull-down experiments or to favour Tau detection with an antibody. We obtained Tau preparations of high purity and yields (between 1 and 3 mg of proteins per 1 l of bacterial culture), suitable for aggregation experiments and other applications.

## RESULTS

### Design of biologically relevant Tau variants

The Tau protein consists of several functionally relevant regions (**Fig. 1A**). The N-terminal region (A2-L243) is important for Tau sorting in the cell and is endowed with a proline-rich domain, whose isomerization is possibly linked to increased aggregation propensity (20). After this stretch, the repeat domains follow (Q244-N368), which consists of fourpseudo-repeats. They are responsible for binding to microtubules and participating to the formation of amyloid structures, in particular the 3^rd^ and 4^th^ repeats (11, 12). The C-terminal region (K369-L441) is physiologically removed when Tau is secreted in the cerebrospinal fluid (21). We aimed to produce three types of Tau constructs: wild type, with pseudo+phosphorylated sites or with increased aggregation propensity.

**Figure 1.**
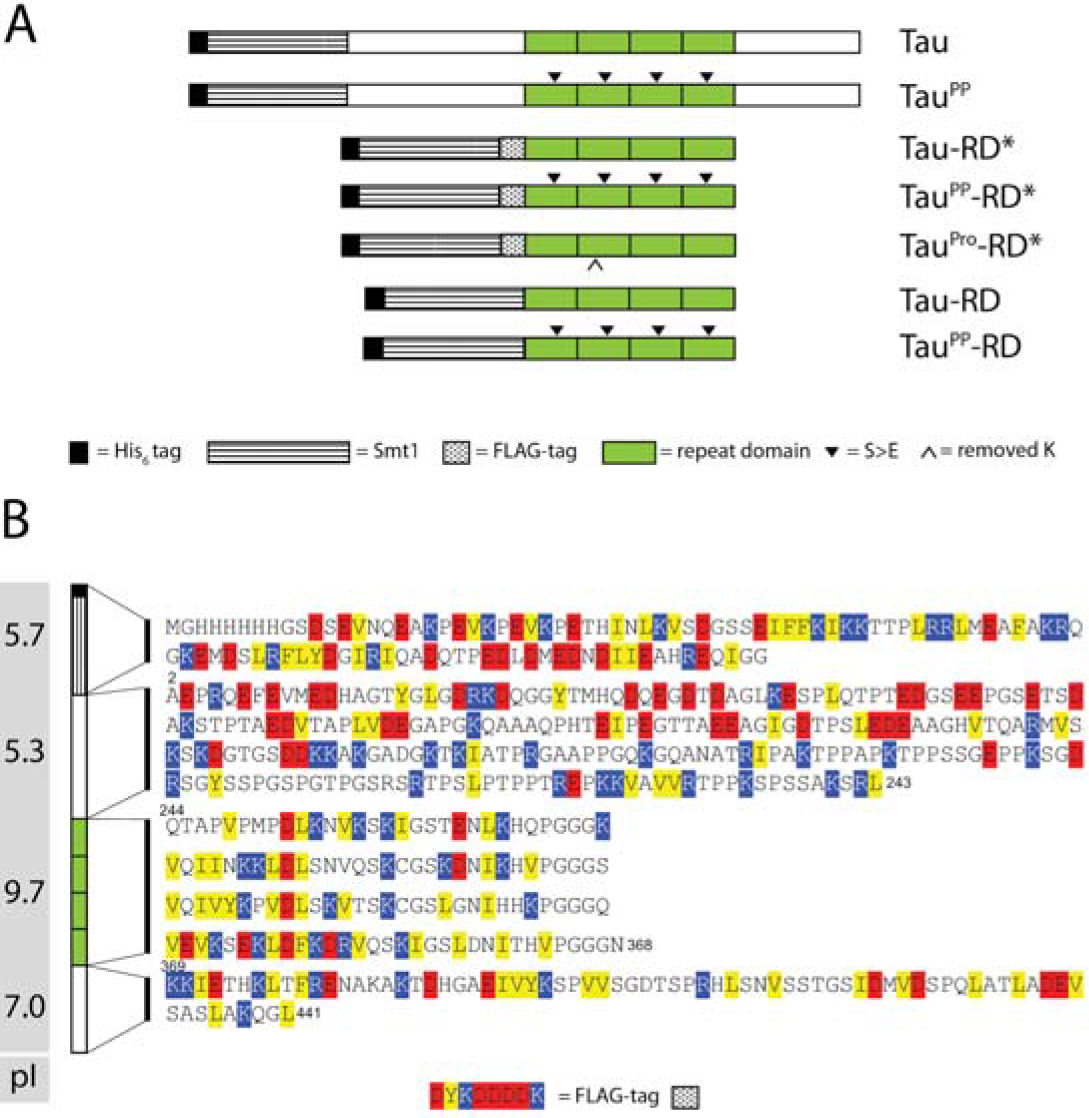
Tau variants used in this study. **a.** Scheme of Tau wild type and Tau variants used in this study. ^PP^: replacement of serines with glutamates at positions 262, 293, 324, 356 (S>E). RD: Repeat Domain. *: FLAG-tag. ^PRO^: Lysine at position 280 removed. **b.** Overview of Tau sequence, hidrophobicity and charge distribution. His_6_-Smt1-Tau sequence colored by hydrophobicity (yellow), negative charges (red) and positive charges (blue). Theoretical Isolectric points (pI) of each domain are shown in the grey box on the left.

A major drawback of studying Tau dynamics is that phosphorylation studies are difficult to perform, as kinases phosphorylase stochastically a subgroup of phosphorylation sites, resulting in heterogenous populations of phospho-Tau species. To this end, we designed a pseudo-phosphorylated Tau construct inspired by previous studies (Tau^pp^, **Fig. 1A**), where replacement of serines with glutamates at defined positions 262, 293, 324, 356 mimics the phosphorylation sites of Tau relevant for disease (22). This approach removes the need for kinases and still offers a functionally relevant protein for studying the impact of negative charges at defined positions.

A second hurdle relates to Tau aggregation dynamics. Indeed, aggregation of full-length Tau takes from several days to up to a week, a time scale where many biological samples become unstable (19). To this end, we generated a truncation of Tau comprising the four repeats (Tau-Q244-E372; Tau Repeat Domain, Tau-RD, also referred to as K18 in the literature ((23)), with reported faster aggregation rates, in the time scale of hours (24). This truncation is an established disease model, causing neurodegeneration in mice (25).

All of our constructs were designed with a cleavable His_6_-Smt tag at the N-terminus. A hexahistidine tag serves as a purification tag, while Smt is a globular domain that favours protein production. After cleavage, Tau protein comprises all residues between A2 and L441 (**Fig. 1B**). Regarding Tau-RD, we added a FLAG tag (DYKDDDDK) at the N-terminus, immediately C-terminal of the cleavage site of Smt1, composed of two Glycines (GG). Thus, the tag is retained after cleavage. It serves for downstream applications, such as western blot detection or pull-down experiments, and positioning at the N-terminus avoids interference with the aggregating regions present toward the C-terminus. Taken together, our designed constructs are suitable to study Tau phosphorylation, aggregation and other aspects of Tau biology.

### Tau sequence properties define strategies for purification

When analysing the amino acidic distribution in Tau, we noted asymmetry in the distribution of charges (**Fig. 1B**). We calculated the isoelectric point (pI) of various segments of Tau based on their composition. While the N- and C-terminal regions (A2-L243 and K369-L441) have a pI of 5.7 and 7.0 respectively, the repeat domain has a pI of 9.7. This implies that at the physiological pH of the cell the N-terminal region is negatively charged, while the repeat domain is positively charged, and the C-terminal region has a near-neutral net charge. Of note, the His_6_-Smt tag has a pI of 5.3, being thus negatively charged at physiological pH. These differentially charged domains can be exploited for binding to different ion exchanges columns, to achieve greater purity of the sample.

### Production and Purification of full-length Tau

First, we aimed to develop a strategy to purify full-length Tau (**Fig. 2A**). First, *E. coli* BL21 DE3 Rosetta 2 cells were transformed with a plasmid encoding for a Tau variant. We chose this cellular strain as it encodes a T7 polymerase necessary to transcribe Tau plasmid. It also encodes for suppressor tRNAs for rare codons, which appear frequently in the human Tau <u>construct. Cells</u> were then induced with Isopropyl β-D-1-thiogalactopyranoside (IPTG) at an optical density (OD) at 600 nm of 0.8, and they were cultured over night at 18°C. Next, they were lysed. Subsequently we heated the lysate at 95°C. This is a key step to increase sample stability as Tau is an intrinsically disordered protein (IDP). Tau cannot, therefore, lose potentially folded structure. The majority of proteins, including proteases, became insoluble and precipitated under these conditions, whereas Tau remained in the supernatant due to its high solubility and lack of folded domains, which could become potentially aggregation-prone after unfolding. Tau-containing supernatant was loaded on an affinity purification column, to which it bound thanks to the His_6_ tag at the N-terminus of the protein. After elution by an imidazole gradient, the protein was further purified with an anion exchange column. Tau was then eluted in a salt gradient. We pooled the Tau containing fractions and cleaved off the His_6_-Smt by adding Ulp1 hydrolase to obtain an untagged Tau species. The resulting protein mix was then loaded on a cation exchange column, where effective separation of Tau from the tag took place upon elution with a salt gradient, due to the different net charges of Tau (positively charged Tau-RD allowed binding to the column) and His_6_-Smt (negatively charged, repelled by the column and found in flow through). This was a critical step to achieve high purity, as differences in the equilibration pH of the cation column yielded less pure eluates, possibly favouring the binding of differentially charged Tau truncation and/or impurities (**Fig. 2B**). While an eluting pH of 7.5 resulted in pure protein, pH of 3. 0, 5.5 and 6.5 yielded less pure protein. We monitored the purification scheme and sample quality by SDS-PAGE (**Fig. 2C**), highlighting the high degree of purity of the final product. We obtained between 5 and 10 mg of proteins per 3.2 l of bacterial culture (n=3).

**Figure 2.**
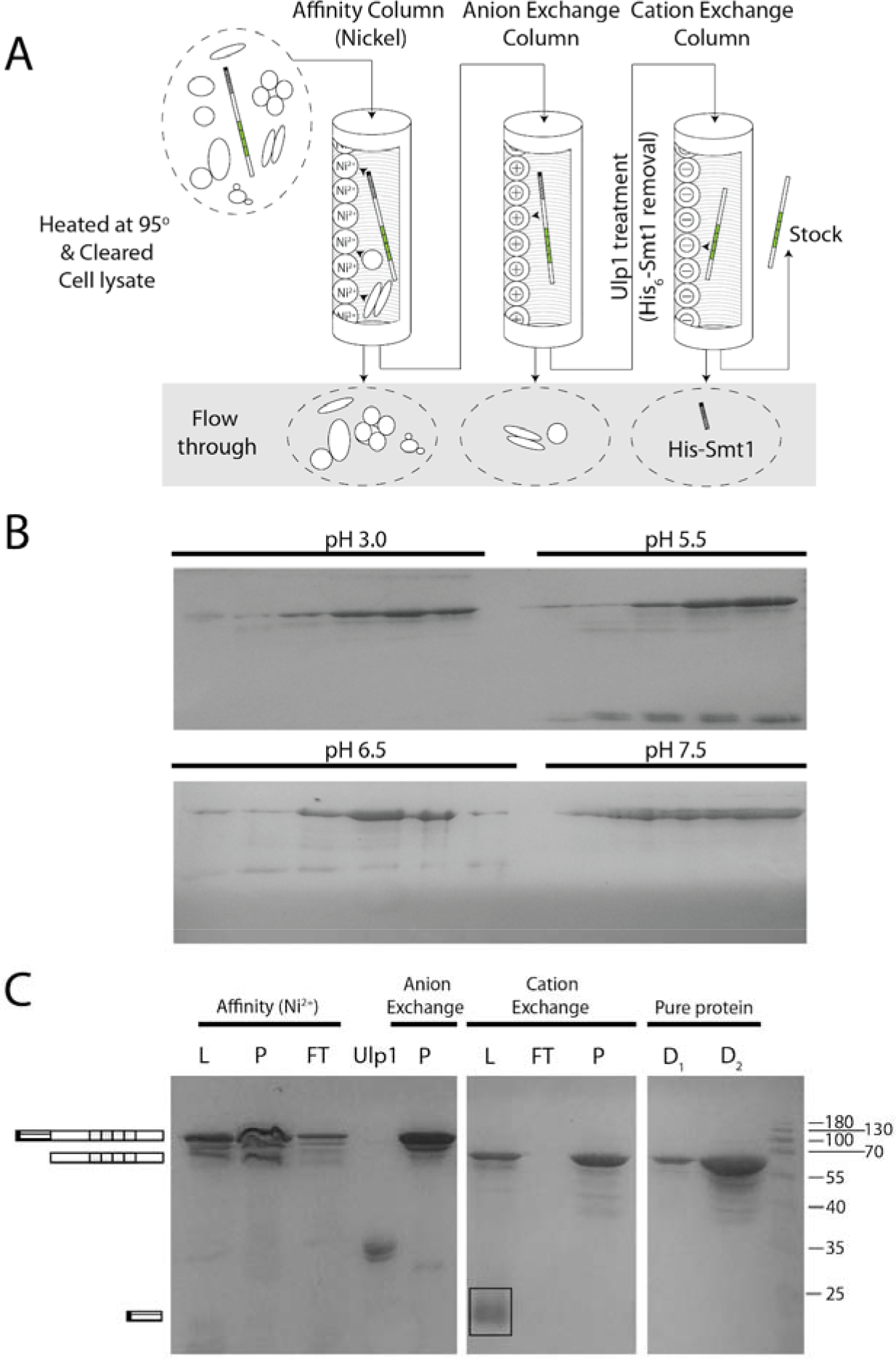
Purification of full-length Tau. **a.** Purification scheme of full-length Tau. **b.** pH scouting to maximize Tau purity. 15% SDS-PAGE gel showing results of pH scouting with cation exchange columns. Four pHs were tested, fractions representative of the whole elution peak were collected and loaded on gel. **c.** Purification of wild type Tau. 15% SDS-PAGE gel showing increased purity of Tau protein as purification proceeds. L: samples loaded on column; P: elution peak; FT: flow through; D1: dilution 1; D2: dilution 2. Boxed band indicates removed SUMO-tag, numbers on the right side of the gel indicate molecular weights in KDa. On the left: schemes of protein segments from Fig. 1A corresponding to different gel bands.

### Robust purification of Tau variants

Next we set out to adapt our protocol for purifying Tau-RD. Remarkably, we could not retrieve protein after heating. We concluded that the highly charged terminal regions increased overall solubility, in line with the observation that Tau-RD is more prone to form fibrils than the full length protein. We consequently adapted the protocol by removing the boiling step (**Fig. 3A**). As Tau-RD has a net positive charge at physiological pH, we also avoided the anion exchange purification step. To compensate for lacking these two steps, we added a size exclusion chromatography step downstream of the purification scheme to remove eventual high molecular weight contaminants. Our adapted protocol resulted in high-purity Tau-RD, as assessed by SDS-PAGE (**Fig. 3B**). We obtained between 4.0 and 8.7 mg of proteins per 3.2 l of bacterial culture (n=5).

**Figure 3.**
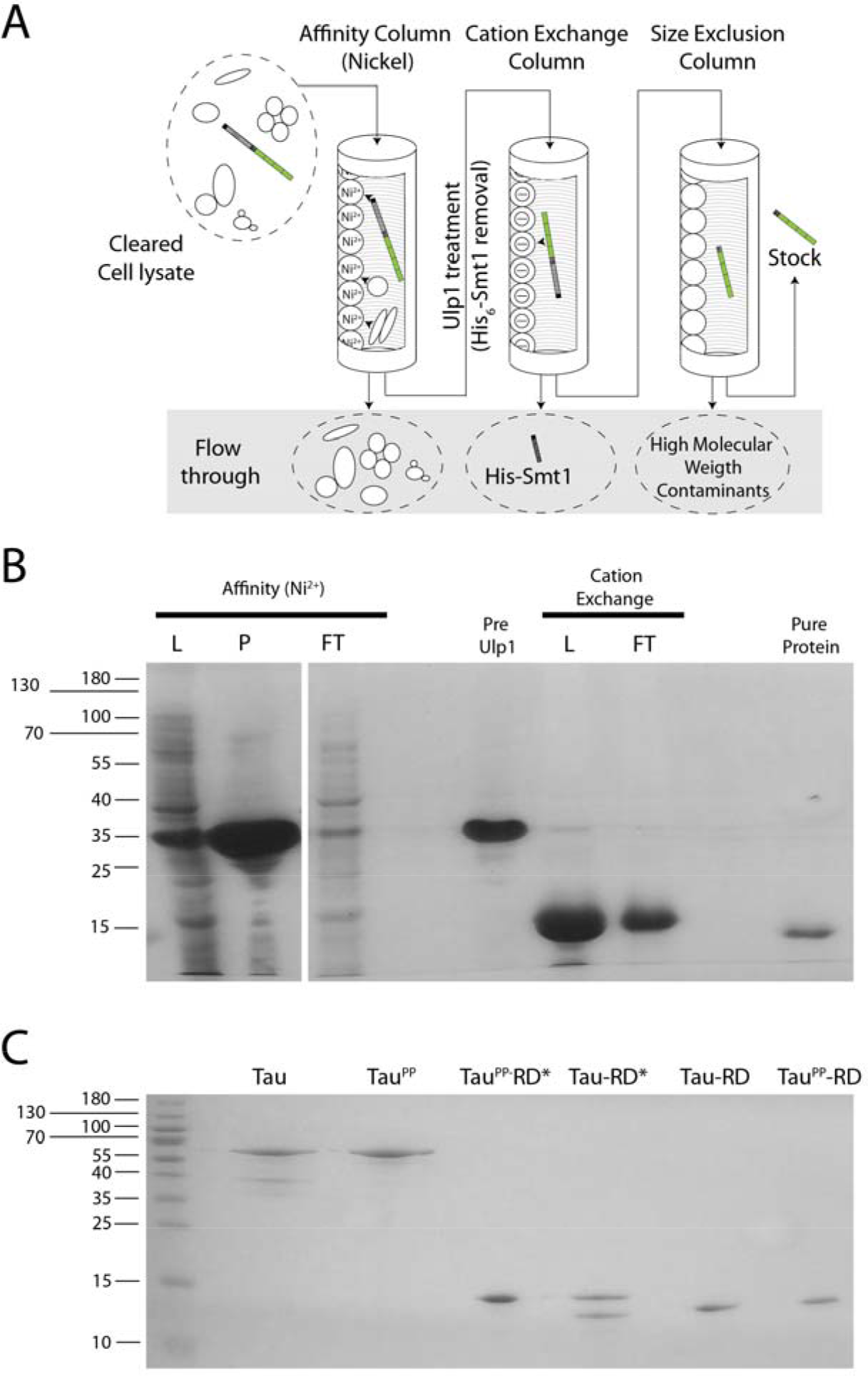
Purification of Tau Repeat Domain. **a.** Purification scheme of Tau-RD. **b.** Purification of wild type Tau-RD*. 12% SDS-PAGE gel showing increased purity of Tau^Pro^-RD* protein as purification proceeds. L: samples loaded on column; P: elution peak; FT: flow through. Numbers on the left side of the gel indicate molecular weights in KDa. **c.** Robust purification of Tau and Tau-RD variants. 15% SDS-PAGE gel showing end products of purifications of different Tau variants. Numbers on the left side of the gel indicate molecular weights in KDa.

Importantly, the two protocols presented for full-length Tau and Tau-RD were extended to other Tau variants, containing for instancepseudo-phosphorylated residues or FLAG-tag, nonetheless yielding pure proteins (**Fig. 3C**). In the case of FLAG-tagged constructs, His_6_-Smt cleavage was extended overnight, as a one-hour treatment was not sufficient to cleave the protein completely. Thus, we could further adapt our protocol developed for full-length Tau to other Tau variants.

### Purified Tau aggregates successfully

We tested the quality of our purified proteins by employing them in an established aggregation assay (26). The negatively charged polymer initiates Tau fibrillation, which is a reliable and reproducible method to study kinetics of Tau aggregation and other aspects linked to Tau-induced neurodegeneration. First, we monitored via SDS-PAGE the aggregation propensity over time of full-length Tau in the presence of increased amounts of heparin (**Fig. 4A**). We observed the appearance of SDS-insoluble aggregates after 6 days of aggregation, at the optimal stoichiometry of four molecules of Tau per molecule of heparin. This is an established stoichiometry to obtain Tau aggregates resembling Alzheimer’s Paired Helical Filaments (27). We did not observe formation of aggregates at sub-stoichiometric Tau:heparin ratios during the time frame of our experiments, whereas excess of heparin interfered with Tau running on the electrophoresis gel, probably due to the high number of negative charges decorating heparin. Thus, our experiment confirmed that our Tau preparation have similar aggregation properties as Tau batches purified with other protocols (18, 19). Compared to other protocols, our purification setup allows purifying Tau variants with different biochemical properties, such as increased aggregation propensity, pseudo-phosphorylated sites or a FLAG-tag at their N-terminus. This versatility does not come at the expense of yields, purity and purification time. Tau preparations were stable even after a week of incubation at 37°C, proving that our purification procedure is highly effective in removing contaminant proteases (**Fig. 4A**).

**Figure 4.**
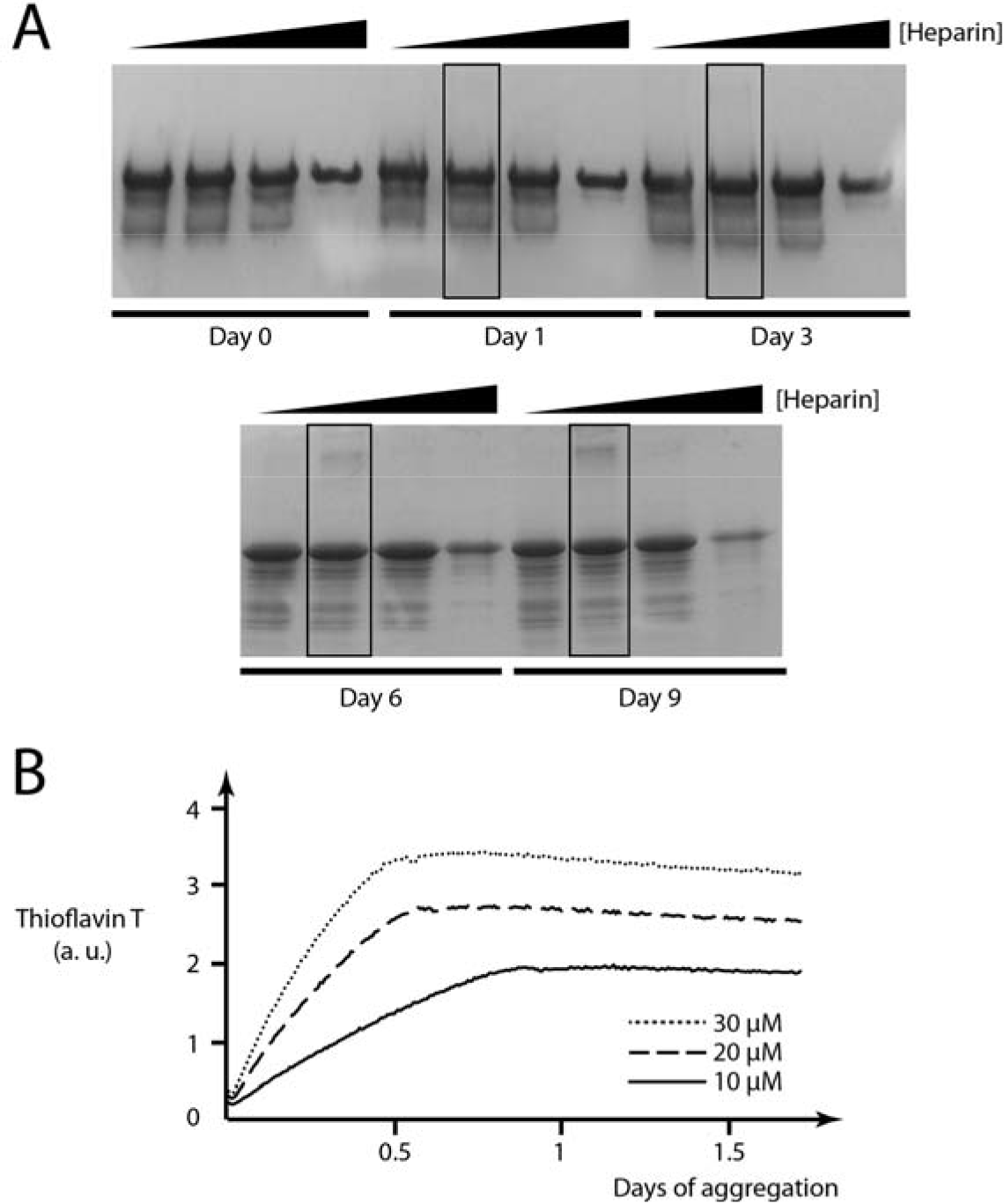
Heparin induces aggregation of Tau variants. **a.** Optimal stoichiometry for heparin-induced Tau aggregation is 1:4 (heparin-Tau). 12% SDS-PAGE gel showing appearance of Tau SDS-resistant protein aggregates over time upon addiction of heparin. Gradients indicate increasing heparin concentrations, boxed lanes indicate optimal heparin:Tau stoichiometry 1:4. **b.** Higher concentration accelerates aggregation of Tau^Pro^-RD*. Thioflavin T assay showing aggregation of Tau^Pro^-RD* at 3 different concentrations (heparin:Tau stoichiometry kept constant at 1:4). Fluorescence correlates with amyloid content and is expressed in arbitrary unit (a. u.).

We further tested the robustness of our protocol by inducing aggregation of Tau-RD domain. We used the Thioflavin T assay to monitor kinetics of aggregation over time. This is an established technique to study aggregation dynamics, in which a fluorescent dye binds to fibrils but not monomers of Tau, resulting in a fluorescent signal allowing monitoring aggregation over time (28). We kept the Tau:heparin ratio as 4:1, this time changing the concentration of monomeric Tau-RD at the beginning of the aggregation, to test (**Fig. 4B**). Increased concentrations of Tau accelerated Tau aggregation, consistent with the proposed model of nucleation-elongation proposed for heparin-induced Tau aggregation (29). Thus, our protocol yields Tau variants whose aggregation dynamics are in agreement with published material.

## DISCUSSION

Here, we presented a protocol to purify Tau species after recombinant production in *Escherichia coli*. Purification of intrinsically disordered proteins is challenging and may require tedious optimization (30). Our protocol allows Tau purification with standard purification equipment and buffers. Compared to previous Tau purification methods, our protocol allows to purify Tau either tag-less or with a tag of choice, thanks to the removable His_6_-Smt tag at the N-terminus of the protein (18, 19). Tau distribution of charges allows effective purification and removal of the His_6_-Smt. Indeed, Tau can both bind to an anion and cation exchange column, the latter used to effectively separate the tag from the purified protein. Most of bacterial proteome has a distribution of isoelectric points lower than 7, thus is negatively charged at physiological pH and removed either by the Nickel column or by the cation exchange column (31). A sub-fraction of the proteome, mostly DNA and RNA binding proteins, is positively charged and thus effectively removed by the anion exchange column. Overall, a double ion-exchange column approach allows effective purification of Tau protein.

Interestingly, the elution buffer of the cation exchange column greatly influences the purity of Tau full-length (**Fig. 2B**). This is probably due to the asymmetric distribution of charges throughout the random coils of Tau protein, exposing different charged patches that bind to the column as pH changes. This distribution is also important for physiological functions of Tau, as the high pI of Tau-RD is necessary for binding to microtubules. In contrast, the N terminal regions of Tau are negatively charged, similar to most cytoplasmic proteins, such that they remain dynamic and exposed even when bound to microtubules (4).

We could not apply the same exact protocol to Tau-RD truncations, as they lack the N-terminus region necessary to bind to the anion exchange column. Tau-RD could not be retrieved after the boiling step, suggesting that the N- and C-termal regions drastically increase Tau solubility. Of note, a study presenting a general purification protocol for disordered proteins based on boiling cell lysates suggests that Tau-RD behaves atypically compared to other disordered segments (32). These differences in solubility may also have had an influence on the evolution of physiologically relevant Tau truncations. Indeed, Tau truncations at the N-terminus are found in humans in physiological conditions, whereas C-terminal truncations were not detected (21). It is tempting to speculate that their higher propensity to aggregate negatively selected them, in favour of more soluble N-terminal truncations.

Our designed Tau constructs, both full-length and RD, aggregate with an optimal Tau:heparin stoichiometry of 4:1, in agreement with published protocols, suggesting that our Tau constructs can be used to model aggregation in Alzheimer and other forms of dementia (28, 29). Indeed, we used N-terminally FLAG-tagged Tau-RD to create Alzheimer-like fibrils, which we could later disaggregate with the human Hsc70 chaperone system (33). To conclude, we present a protocol yielding pure and stable Tau constructs suitable for a wide array of *in vitro* studies.

## ACKNOWLEDGMENTS

We are grateful to Ineke Braakman for continuous support. We thank Madelon Maurice for collaboration in the Initial Training Network “WntsApp” (No. 608180)], supported by Marie-Curie Actions of the 7th Framework programme of the EU. SGDR was further supported by the Internationale Stichting Alzheimer Onderzoek (ISAO; project “Chaperoning Tau Aggregation”; No. 14542) and a ZonMW TOP grant (“Chaperoning Axonal Transport in neurodegenerative disease”; No. 91215084).

## AUTHOR CONTRIBUTIONS

S.G.D.R. and L.F. conceived the study; S.G.D.R., L.F. and F.G.F. planned experiments; L.F., W.J.C.G., M.v.W., R.K., K.K. and L.v.B. did experiments; L.F., W.J.C.G., M.v.W., R.K., K.K. and L.v.B. analysed data; S.G.D.R. and L.F. wrote the manuscript, with contributions of all authors.

## COMPETING INTERESTS

The authors declare no competing interests.

## MATERIALS AND METHODS

### Isoelectric points calculation

Theoretical isoelectric points were calculated using ProtParam tool (https://web.expasy.org/protparam/).

### Purification of Tau variants

We designed two types of Tau constructs, full-length and Repeat Domain (**Fig. 1A**). We propose two purification protocols, based on the variants to be purified.

### Purification of human full-length Tau

Following, a step-by step procedure to purify human full-length Tau:

1. Human Tau and Tau^PP^ were overproduced recombinantly in *E. coli* BL21 Rosetta 2 (Novagen). Cells were induced with 0,15 mM Isopropyl β-D-1-thiogalactopyranoside (IPTG, Sigma) at Optical Density at 600 nm (OD600) of 0.8 and cultured over night at 18°C.
2. Cells were harvested, flash-frozen in liquid nitrogen and stored at −80°C until further usage.
3. Pellets were thawed in a water bath at 37°C and resuspended in 50 mM HEPES-KOH pH 8.5 (Sigma-Aldrich), 50 mM KCl (Sigma-Aldrich), ½ tablet/50 ml EDTA-free protease inhibitor (Roche), 5 mM β-mercaptoethanol (Sigma-Aldrich).
4. Cells were disrupted by an EmulsiFlex-C5 cell disruptor (Avestin).
5. Lysate was cleared by centrifugation, filtered with a 0.22 μm polypropylene filter (VWR) and supernatant was purified using an ÄKTA Purifier chromatography system (GE Healthcare).
6. Lysate was loaded onto a POROS 20MC (Thermo Fischer Scientific) affinity purification column in 50 mM HEPES-KOH pH 8.5, 50 mM KCl, 5 mM β mercaptoethanol, eluted with a 0-100% linear gradient (5 Column Volumes, CV) of 0.5 M imidazole.
7. Fractions of interest were collected, diluted to salt concentration <100 mM and loaded onto a POROS 20HQ (Thermo Fischer Scientific) anion exchange column equilibrated with 50 mM HEPES-KOH pH 8.5. Protein was eluted with a 0-100% linear gradient (15 CV) of 1 M KCl (Carl Roth).
8. The His_6_-Smt tag was removed by Ulp1 treatment (purified in house), shaking at 4°C for 30 minutes.
9. Protein was diluted to salt concentration <100 mM and loaded onto a POROS 20HS (Thermo Fischer Scientific) cation exchange column equilibrated with 50 mM HEPES-KOH pH 7.5. Protein was eluted with a 0-100% linear gradient (15 CV) of 1 M KCl (Carl Roth).
10. Fractions of interest were further concentrated to the desired final concentration using a concentrator (Vivaspin, cut-off 5 kDa).
11. Protein concentration was determined using an ND-1000 UV/Vis spectrophotometer (Nanodrop Technologies) based on the theoretical extinction coefficient at 280 nm and purity was assessed by SDS-PAGE. Protein was aliquoted and stored at −80°C.

### Purification of human Tau-RD

Following, a step-by step procedure to purify human Tau-RD:

1. Human Tau-RD*, Tau^PP^-RD*, Taupro-RD*, Tau-RD, Tau^PP^-RD were overproduced recombinantly in *E. coli* BL21 Rosetta 2 (Novagen). Cells were induced with 0,15 mM IPTG (Sigma) at Optical Density of 0.8 and cultured over night at 18°C.
2. Cells were harvested, flash-frozen in liquid nitrogen and stored at −80°C until further usage.
3. Pellets were thawed in a water bath at 37°C and resuspended in 50 mM HEPES-KOH pH 8.5 (Sigma-Aldrich), 50 mM KCl (Sigma-Aldrich), ½ tablet/50 ml EDTA-free protease inhibitor (Roche), 5 mM β-mercaptoethanol (Sigma-Aldrich).
4. Cells were disrupted by an EmulsiFlex-C5 cell disruptor (Avestin).
5. Lysate was cleared by centrifugation, filtered with a 0.22 μm polypropylene filter (VWR) and supernatant was purified using an ÄKTA Purifier chromatography system (GE Healthcare).
6. Lysate was loaded onto a POROS 20MC (Thermo Fischer Scientific) affinity purification column in 50 mM HEPES-KOH pH 8.5, 50 mM KCl, 5 mM -β mercaptoethanol, eluted with a 0-100% linear gradient (5 Column Volumes, CV) of 0.5 M imidazole.
7. Fractions of interest were collected and concentrated in a buffer concentrator (Vivaspin, cut-off 10 kDa) to final volume of 3 ml.
8. The concentrated sample was desalted with a PD-10 desalting column (GH Healthcare) in 50 mM HEPES-KOH pH 8.5, ½ tablet/50 ml Complete protease inhibitor (Roche)) and 5 mM β-mercaptoethanol.
9. The His_6_-Smt tag was removed by Ulp1 treatment, shaking at 4°C for 30 minutes (increased to 16 hours when purifying FLAG-containing Tau variants).
10. Next day, protein was diluted to salt concentration <100 mM and loaded onto a POROS 20HS (Thermo Fischer Scientific) cation exchange column equilibrated with 50 mM HEPES-KOH pH 8.5. Protein was eluted with a 0-100% linear gradient (15 CV) of 1 M KCl (Carl Roth).
11. Fractions of interest were collected and loaded onto a HiLoad 26/60 Superdex 200 pg (GE Healthcare Life Sciences) size exclusion column equilibrated with aggregation buffer (25 mM HEPES-KOH pH 7.5, Complete Protease Inhibitors (1/2 tablet/50 ml), 75 mM KCl, 75 mM NaCl and 10 mM DTT).
12. Fractions of interest were further concentrated to the desired final concentration using a concentrator (Vivaspin, cut-off 5 kDa).
13. Protein concentration was determined using an ND-1000 UV/Vis spectrophotometer (Nanodrop Technologies) based on the theoretical extinction coefficient at 280 nm and purity was assessed by SDS-PAGE. Protein was aliquoted and stored at −80°C.

### Sodium Dodecyl Sulfate - Polyacrylamide Gel Electrophoresis (SDS-PAGE)

20 μL of samples were collected for each purification/aggregation step of interest and were supplemented with 20 μL of 2x sample buffer (0.625 M Tris pH 6.8 (Sigma-Aldrich), 12.5% glycerol (CARL ROTH), 1% SDS (Bio-Rad), 0.005% Bromophenol Blue (Bio-Rad), 5mM freshly added β-mercaptoethanol (Sigma-Aldrich). 15% SDS gels were prepared (Separation buffer: 0.38 M Tris pH 8.8, 15% acrylamide(National Diagnostics), 0.1% SDS, 0.1% APS (Sigma-Aldrich), 0.04% TEMED (Sigma-Aldrich), Stacking buffer: 0.125 M Tris pH 6.8, 4% acrylamide, 0.1% SDS, 0.075% APS, 0.1% TEMED) and run in 1x Laemmli buffer (0.025 M Tris base pH 6.8, 0.152 M glycine (SERVA Electrophoresis GmbH), 0.1% SDS, diluted from 10x stock). Gels were stained with Coomassie staining solution (0.2% Coomassie Brilliant Blue (SERVA Electrophoresis GmbH), 45% methanol (Interchema Antonides-Interchema), 10% acetic acid (Biosolve) and 55% demineralised water) and destained using destaining solution (30% methanol, 10% acetic acid and 60% demineralised water).

### Formation of Tau fibrils

Monomeric human full-length Tau was aggregated by adapting a published procedure (34). Briefly, 50 μM of Tau were mixed with varying concentrations of heparin (0.125, 1.25, 12.5 and 125 μM) in 25 mM HEPES-KOH pH 7.5, 75 mM KCl, 75 mM NaCl, 1 mM DTT and Heparin Low Molecular Weight (Santa Cruz Biotech). Samples were heated at 95°C for 10’ and supplemented with Complete Protease Inhibitors (1/2 tablet/50 ml) and 1 mM DTT. Aggregation was performed at 37°C for shaking in an incubator at 180 rpm. Samples were collected at time points described in **Fig. 4A**.

Monomeric human N-terminally FLAG-tagged Tau-RD was aggregated in 25 mM HEPES-KOH pH 7.5, Complete Protease Inhibitors (1/2 tablet/50 ml), 75 mM KCl, 75 mM NaCl, 10 mM DTT and Heparin Low Molecular Weight (Santa Cruz Biotech), concentrations depending on the experiment and Tau:heparin ratio were kept at 4:1. Aggregation was performed at 37°C, shaking at 180 rpm, and aliquots were flash frozen at time points indicated in the text. Aliquots were then thawed in a water bath at 37μC for downstream applications.

### ThioflavinT assay

N-terminally FLAG-tagged Tau-RD (100 μl) of varying concentrations, 60 μM ThioflavinT (Sigma) and 5μM heparin Low Molecular Weight were aggregated in aggregation buffer in a transparent, lidded Greiner 96-well plate (Sigma-Aldrich). Fluorescent spectra were recorded with a SpectraMax i3 (Molecular Devices) every 10 minutes for 16 hours.

